# Dynamic Order in Allosteric Interactions

**DOI:** 10.1101/2020.01.27.920850

**Authors:** Sina Türeli, Türkan Haliloğlu

## Abstract

Allostery is an intrinsic dynamic phenomenon that underlies functional long-distance interactions in proteins, which we study here by stochastic calculus approach to elastic network models (ENMs). We show that once you drop the usually accepted high friction limit and include hydrodynamic interactions in ENMs, a simple measure that uses the pairwise difference in the time-delayed correlations of residue fluctuations provides insight about functional sites and their dynamical behaviour in allosteric communication. We present this with three exemplary cases Aspartate Carbamoyl transferase, Insulin Receptor and DNA-dependent Protein Kinase. We show that proteins possess characteristic pathways operating at different time-delay windows with slow to faster motions underlying the protein function. As these pathways help communication between key residues of functionality, they can also be used to identify their locations without any prior knowledge other than the protein crystal structure.

## 1 Introduction

Allostery is a fundamental mechanism through which protein activity is regulated. When cellular conditions such as pH or the concentration of certain molecules change, proteins respond by regulating their activities such as signalling and enzymatic function. In the simplest case, a protein has a *regulator site* (a turn on/off site) which is altered by the change of conditions and a *regulated site* which is affected by the change in the regulator site. A change in the regulated site usually causes a modification in function. Proteins adopt to the needs of the cell through such *allosteric sites*. Although the underlying mechanism is still being debated, it is generally accepted that allosteric interactions appear with subtle thermodynamic and dynamical changes [10].

One of the great challenges of relating protein dynamics to functionality is to determine the allosteric sites and how they communicate in relation to the protein function. Optimized contact topologies and dynamics provide a framework at which this communication is efficient. Elastic Network Models (ENMs) have found great use to disclose functional behaviour in relation to its equilibrium dynamics versus contact topology in the native state [30]. Two of the most popular ENMs used for this purpose are the Anisotropic Network Model [2] (ANM) and the Gaussian Network Model [14] (GNM), both of which assume the high friction regime.

In the assumed ensemble of conformations, ENMs predict to what extent fluctuations of residues *I* and *J* are correlated. More preciselyif Δ*R*_*I*_ (*t*) and Δ*R*_*J*_ (*t*) are the fluctuationsat time *t* of residues *I* and *J* from their initial position,then the normalized cross-correlation between residues *I* and *J* is given by

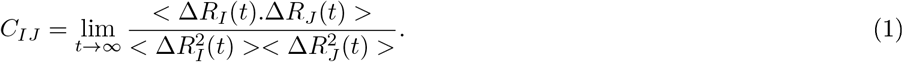

The ENMs correlation between two residues’ fluctuations decay exponentially with distance and are symmetric with respect to residue pairs. Due to the decay one cannot score the strength of communication based on the strength of correlations alone and needs to introduce additional complicated statistical approaches to factor out the distance effect (see [1]) or have some a priori knowledge of the allosteric sites to predict the rest (see [1, 11, 24]). The symmetry with respect to indices mean that one cannot study the “topology of communication” using *C*_*IJ*_ – *C*_*JI*_ neither.

However, ideally, one should be able to infer the communication pathways simply by studying the connection topology of the protein. Starting with [12], this idea have been given a more formal and elegant perspective by employing tools from information theory [26] (see [21, 19, 13] for non ENM versions). These methods require the use of normalized time delayed cross-correlations defined as

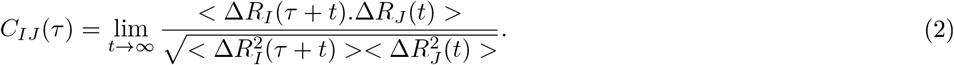

This gives information about how much the fluctuations of residue *J* at some time affects the fluctuations of residue *I* some *τ* time later. One can expect that this should contain more information about allosteric behaviour than equal time correlations, since allostery is a series of chronological events rather than simultaneous. In principle, information about causality due to geometric organization of residues should already be contained in Δ|*C*_*IJ*_ (*τ*)| = |*C*_*IJ*_(*τ*)| − |*C*_*IJ*_(*τ*)| ([12]). For instance, if there is a *J* such that Δ |*C*_*IJ*_ (*τ*)| > 0 for a large group of other residues *I*_1_, …, *I*_*n*_, it is expected that *J* should have some functional importance. This means that *J* leads and possibly determines the dynamics of the other parts of the protein and not the other way around.

However, in high friction regime, Δ |*C*_*IJ*_ (*τ*)| is symmetric with respect to *I* and *J* and cannot be used on its own for studying directionality. As far as we are aware, in all such allostery studies where an ENM is used, the high friction assumption is made. That is why it is either one needs to use more complicated quantities such as transfer entropy or more indirect methods as highlighted above. One of the advantages of our method is that Δ |*C*_*IJ*_ (*τ*)| is generally non-zero but is still an ENM and therefore fast. In this way, we use |Δ *C*_*IJ*_ (*τ*)| to find preceding and lagging residues in a protein and demonstrate that these are related to allostery. Some of the other main advantages of this method is that one does not require a priori knowledge of regulator or regulated sites, no statistical scoring is required. This model also integrates a “time scale” into the model which allows us to identify allosteric patterns that emerge from both slow and fast motions (see for instance [4]). To test our model DORIA (short for Dynamical Order in Allostery), we consider three cases, a text book example Aspartate Carbamoyltransferase (ATCase) and two more contemporaray examples, Insulin Receptor (IR) and DNA-dependent Protein Kinase (DNA-pkc).

## 2 Methods

The slow modes of an ENM represent large domain movements (Fig 1A) and few slow modes are usually sufficient to understand the main functional moves of the protein [3]. In this sense they are macroscopic from the point of view of the cell since they can impart work on the cell by cutting, pushing etc. It has also been demonstrated in many cases that slow modes are related to the motion a protein exhibits when it switches from apo to holo conformation or vice versa upon allosteric activation [15, 23]. The set of modes after the few first slowest should become important when one wants to understand how smaller parts of the protein communicate with each other (these modes as microscopic from the point of view of the cell but macroscopic from the point of view of the protein). It has indeed been demonstrated in theoretical works that a higher number of modes becomes important in allosteric communication and that elastic networks can be employed successfully for prediction of functional sites from the knowledge of an allosteric site ([11]). It has also been demonstrated in experimental studies that even in proteins where the response to allosteric activators is usually carried out by slow modes, intrinsic rearrangement of faster modes can also drive allosteric activation [31, 32]. Therefore when studying allostery a method that spans a wider range of modes is essential.

**Figure 1:**
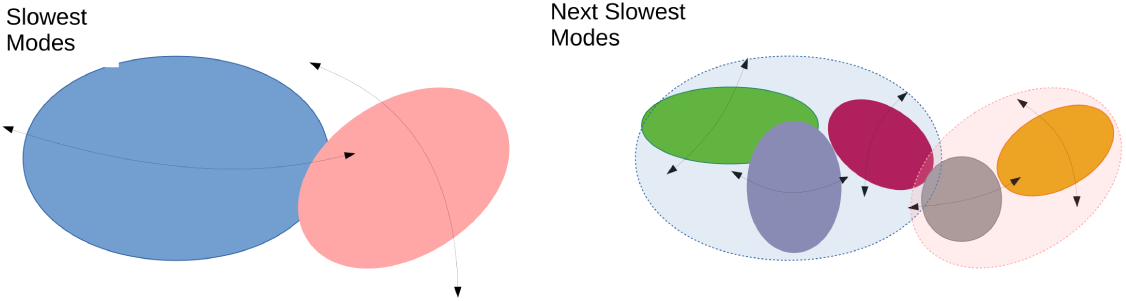
Diagram on the left represents hypothetical slowest modes in which larger domains move like rigid bodies and diagram on the right represents next set of slowest modes in which are faster and smaller domains oscillate as smaller sub-dynamic domains.

It is expected that if a residue *I* has the right connections then given enough time it can affect the behaviour of a residue *J* by energy transfer. However, it might be the case that the other direction of energy flow is not so efficient (i.e the process that leads to energy flow *I* → *J* is mostly irreversible). Then Δ |*C*_*IJ*_ (*τ*)| − Δ |*C*_*JI*_ (*τ*)| should be non-zero for some *τ*. Moreover, it is reasonable to expect that patterns for shorter delay represent interactions due to slowest modes whereas those for longer delay following set of modes (the reasoning behind this is explained in more detail in the supporting information). This idea forms the basis of our approach in which we find residues whose dynamics generally *precedes* the dynamics of other parts of the protein and reversely those which generally *lag*.

To find the preceding and lagging residues more efficiently we use a quantity called the mutual information which is a simple generelizationof delayedcross-correlation(see SI).Ve denote the mutual information between residues *I* and *J* with delay *τ* as *M*_*IJ*_ (*τ*). Using this,we constructsome measuresof communicationbetweenresidues:

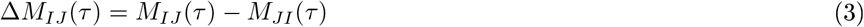

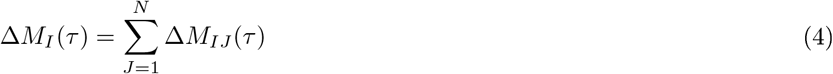

We call the first object the *communication profile*. Positive values for Δ*M*_*IJ*_ (*τ*) imply that *J* precedes the dynamics of *I*. In high friction ENMs both *M*_*IJ*_ (*τ*) and *C*_*IJ*_ (*τ*) are symmetric with respect to *I*,*J* and therefore Δ*M*_*I*_ (*τ*) = 0. Here we do not employ the high friction limit. The way to build these models as explained in SI. This allows us to study communication profiles in ENMs. We call the second object the *mean communication profile*. Positive value for this means that the residue *J* in overall precedes the dynamics of the rest of the protein while negative value means that it in overall this residue lags. Zero values mean that it precedes the dynamics of some of the regions where as it lags with respect to some others. An important parameter in these two objects is the delay *τ*. Our analysis has revealed that there are some very apparent patterns that appear in short and long delay times. Determination of these, respectively called the *short time delay scale* and *long time delay scale* are also described in SI. Motivated by the results we observe; we refer to the short delay patterns as the *global-dynamic motions* whereas the long delay patterns as the *sub-dynamic motions*. This also has the reason that slow mode movements transmit their effect from one end to another in a much shorter delay (since they essentially move like a rigid body) but less slow mode movements involve more sub-dynamic components interacting with each other and take a longer time to transmit.

## 3 Results

### 3.1 ATCase

ATCase catalyzes the reaction of L-aspartate (L-a) and carbomoyl phosphate (CP) to N-carbomoyl-L-aspartate (CAA). It is the first step of the pyrimidine biosynthetic pathway whose end products are Cytidine triphosphate (CTP) and uridine triphosphate (UTP). ATCase is a hexamer and each monomer is made of a catalytic domain (C domain) and a regulatory domain (R domain). The C domain binds the substrates L-a and CP, whereas the R domain binds molecules CTP and UTP (inhibitors) or ATP and GTP (activators).

ATCase is known to have two states; the tense state (T-state, 2ATC) and the relaxed state (R-state, 2AT1), which co-exist in equilibrium. The T-and R-states have respectively low and high affinity for substrates. There is a rotation in the R domains between the T-and R-states (see Fig 2). When the substrates bind to the catalytic sites of the T-state, this is followed by an immediate conformational change to the R-state [22, 29, 33, 20, 5]. In the R-state, the catalytic sites move closer and facilitate the reaction while the catalytic cavity is enlarged, resulting in higher affinity. When substrates are abundant, ATCase stays in the R-state and the substrates at ATCase are converted to CAA. When the end products CTP and UTP bind to the R domain, they stabilize the T state by curling the N-terminal inwards. On the other hand, when ATP and GTP, the end products of another parallel biochemical pathway, bind to the R domain, they push the N-terminal in a way that stabilizes the interface and locks the ATCase in the R-state. The molecules that bind to the R domain, however, do not cause immediate large conformational changes but alters the energy landscape in a way in which one state is preferred over the other state. We refer to the sites where the substrates or their analogues bind as catalytic sites and where the regulating molecules bind as regulator sites.

**Figure 2:**
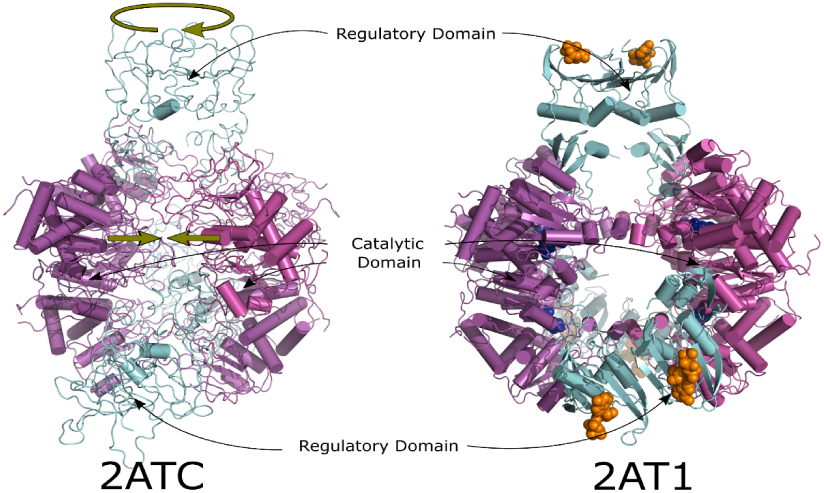
ATPCase in the T-state (2ATC) and R-state (2AT1) of the C and R domains colored in magenta and cyan, respectively. Blue sticks are the substrates (PAL in this case) near the catalytic site and orange ones display the molecules that bind to the R domain (in this case ATP). The positions of the substrates and regulator molecules are from 4KGV for the demonstration (not included in the calculations). In the transition from the T-state to the R-state, the R domains rotate around the symmetry axis of the protein and force the C domains to close the gap in between (the green arrows).

In this case, we run DORIA for the two conformations of ATCase: 2ATC (the T-state) and 2AT1 (the R-state).

#### 3.1.1 Short time delay scale

T-state (2ATC): Most of the positive peaks are concentrated at the C domain at/near the active site (see Fig 3 second row). There are also some positive peaks in the R domain, shorter than most of the peaks at the C domain. All of the negative peaks are at the R domain with the tallest (highest negative) points of the peak usually being at the regulator sites.

R-state (2AT1): The distribution of the positive and negative peaks is very similar to 2ATC (see Fig 3 second row). However, there is a subtle change that might be of significance. The relatively short positive peak near Asn234-Leu240 in 2ATC becomes the dominant positive peak at the C domain. It is likely that this stretch of residues is important for the conformational transition from the R to T-state.

**Figure 3:**
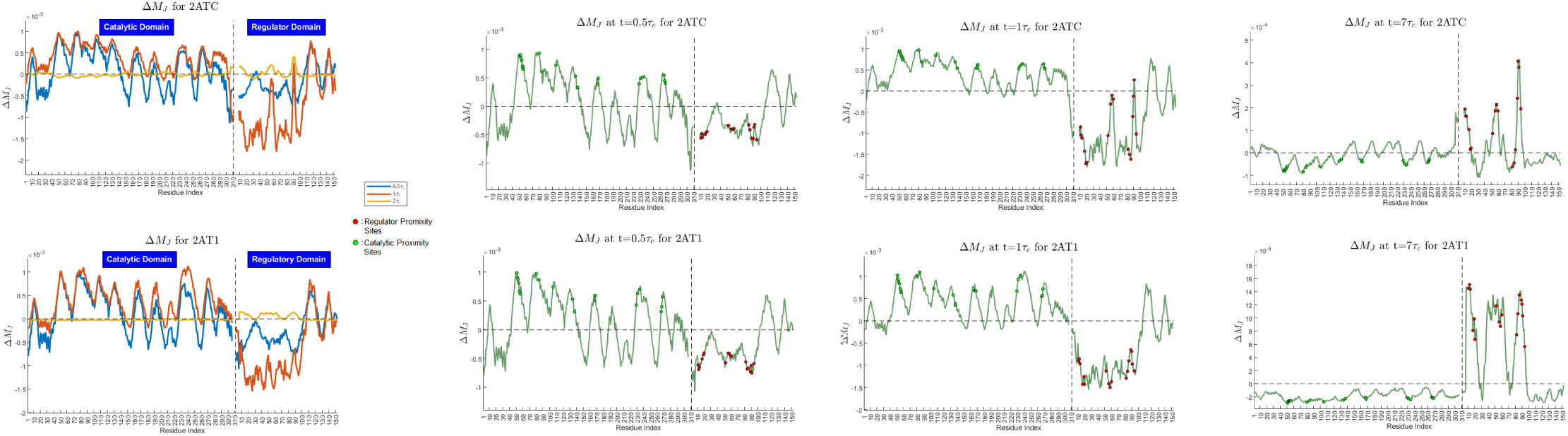
Δ*M*_*J*_ of 2ATC and 2AT1 at the short, normal and long time delay scales. The disks indicate functionally important residues whose distance to the substrates and regulator molecules or substrates (calculated from the PDB file 4KGV) are less than/equal to 5 *A*°, which we call catalytic and regulator proximity sites.

**Figure 4:**
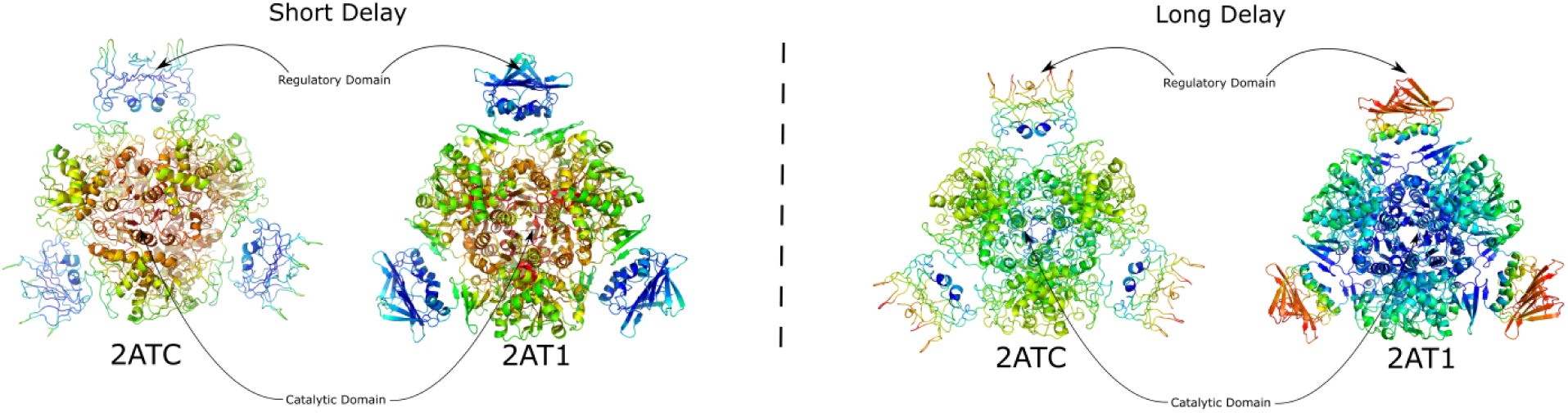
ATPCase in the inactive T-state (2ATC) and the active R-state (2AT1) colored according to Δ*M*_*J*_ values at the short and long-time delay scales (negative values linearly to blue-cyan-green and positive values linearly to green-yellow-red color scales).

The communication profile plots in Fig S1 reveal that for both cases the regions with positive peaks precedes the dynamics of regions with negative peaks. All thus imply that the C domain (active sites) plausibly determines the dynamics of the R domain (regulator sites) at shorter time delay scales. This observation is in accordance with the experimental findings that the substrate binding to T-state results in conformational changes to the R-state [20]. However, since this relatio also exists in the R-state, it is possible that changes near the active site (such as conversion of substrates to products and their release from the cavity) could also be a driving factor for the opposite transition. Note that this finding also supports the idea that shorter time delays represent the effect of more global domain movements (which are also in the spectrum of slow dynamics) through cooperativity of rigid domains.

#### 3.1.2 Long time delay scale

T-state (2ATC): The positions of the negative and positive peaks are exchanged compared to the short time delay scale (see Fig 3 fourth row). In this case, we see very high positive peaks at the R domain with the tops at the regulator sites. There are also some short positive peaks at the C domain that correspond to the regions closer to the R domain. On the other hand, we see that there is a distribution of shorter negative peaks at the C domain with the catalytic sites at the top of the peaks. There are also a few negative peaks with similar height at the interface of the R and C domains. This corresponds to the neck that connects the two domains.

R-state (2AT1): The distribution of the peaks is like the T-state, yet the peaks are much more emphasized (see Fig 3 fourth row). The positive peaks at the R domain are much taller and the C domain has only negative peaks. This displays the intrinsic behavior of dynamic order in the allosteric communication in the T-state.

In the 2D communication profile plots in Fig S1, we clearly see the effect of the R domain on the rest of the protein in both states, being more emphasized in the R-state. This is likely because the subdomains of the R domain form an interface in the R-state and this cluster of sub-domains act as a ‘soft hinge’ whose dynamics affect the dynamics of the rest of the protein via a larger group of slow to medium modes. This pathway inheritably exists in the T-state. We thus suggest that the R domain (regulator sites) regulates the behavior of the C domain (active sites) via sub-dynamic cooperative motions at longer time delay scales with the regulatory behavior much more emphasized in the relaxed state. This is also supported by experimental evidence that the regulatory molecules enforce gradual conformational change by either stabilizing or destabilizing the R-state [33, 20, 5].

### 3.2 Insulin Receptor (IR)

The IR is a dimeric transmembrane protein activated by insulin. Its domains starting from the extra-cellular side are in order: Leucine-rich 1 (L1), Cysteine-rich (CR), Leucine-rich 2 (L2), fibronectin type III 1, 2 (FNIII-1, FNIII-2) and insertion *α* (ID-*α*) domains, *α*-CT helix, fibronectin type III 3 (FNIII-3) and insertion *β* (ID-*β*) domains, transmembrane helix (TM) and Tyrosine-Kinase (TK) domain. The TM connects the extra-cellular part to the intra-cellular TK domain. The L1, CR, and L2 domains are respectively connected by the linkers 1 and 2. The classically known binding groove for the insulin is formed by the L1 domain, the *α*-CT helix and the FNIII-1 domain (binding sites 1 and 1’). Another suggested insulin binding location resides between the FNIII-1 and FNIII-2 domains (binding sites 2 and 2’). It is suggested that the binding sites 1/2’ have a negative allosteric effect on the binding sites 1’/2 [34, 7]. With the insulin binding to 1, the structural changes in the FNIII domains is assumed to transduce the “change of state” signal. This activates the TK domain [25], in which the L1, CR, and L2 domains retract upwards and bend inwards toward the FNIII-1 domain (see Fig 5). This is obtained by the rotations of the L1-CR-L2 domains around the linkers 1 and 2. *α*-CT helix moves about 50 *A*°, which in some sense acts like a ligand and is embedded between the L1-L2-FNIII-1 domains in the holo state. On the other hand, glycogens are bound to the extracellular part at various locations (in both states, in the same positions).

**Figure 5:**
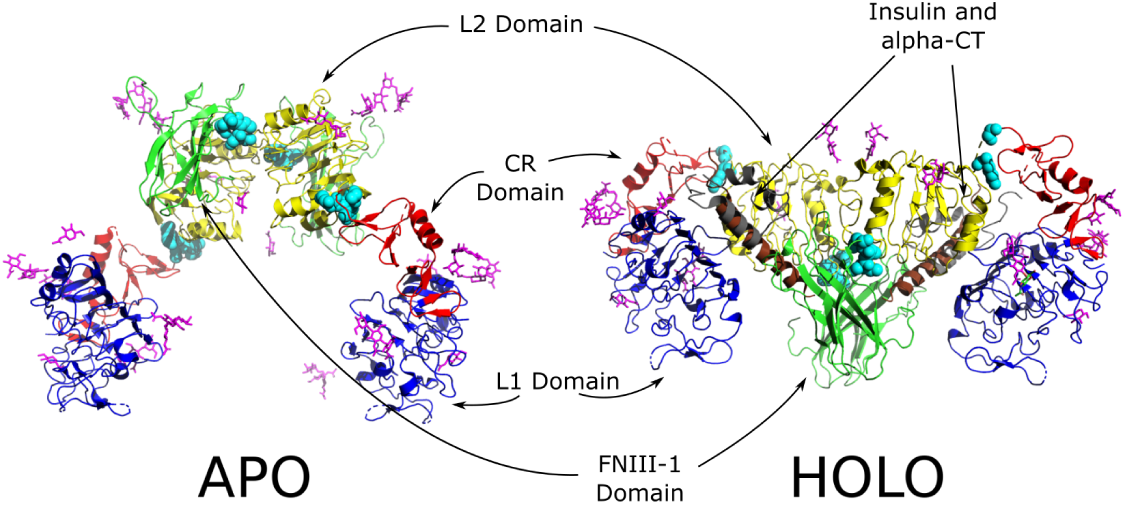
The IR in the apo (4ZXB, X-Ray 3.3*A*°) and holo (6CE9, Cryo-EM 4.8 *A*°) states. The L1, CR, L2 and FINIII-1 domains are shown in blue, red, yellow and green, respectively. Cyan balls represent the linkers 1 and 2, the insulin molecules are in grey and the *α*-CT helices are in brown (shown in the holo state). Magenta sticks are glycogens that bind to the IR in both states.

We run DORIA on the IR at both apo and holo states. The apo and holo states contain, respectively, glycogens only and glycogens and insulin molecules. The alpha-CT helices exist in both. There is also a cryo-EM apo form (with lower resolution), which seems to differ from 4ZXB structure. It is thus possible that 4ZXB might be a sample from the apo state dynamic ensemble [25, 6]. Moreover, the holo state contains the domains only up to FNIII-1 plus the alpha-CT helix, while the apo state is up to and including the FNIII-3 domain. We have thus not included the FNIII-2 and FNIII-3 domains to be able to compare the apo and holo states.

#### 3.2.1 Short time delay scale

In the apo state, most of the highest positive peaks appear at or near the *α*-CT helix binding sites (which also includes the insulin molecule proximity sites) within the L1 and L2 domains and at the glycogen proximity sites of the L1 domain (Fig 6). The tallest negative peaks are concentrated at the linker 1 including the second half of the L1 and CR domains. Some of the glycogen proximity sites near the C-terminal of the L1 domain and the CR domain also display negative peaks.

**Figure 6:**
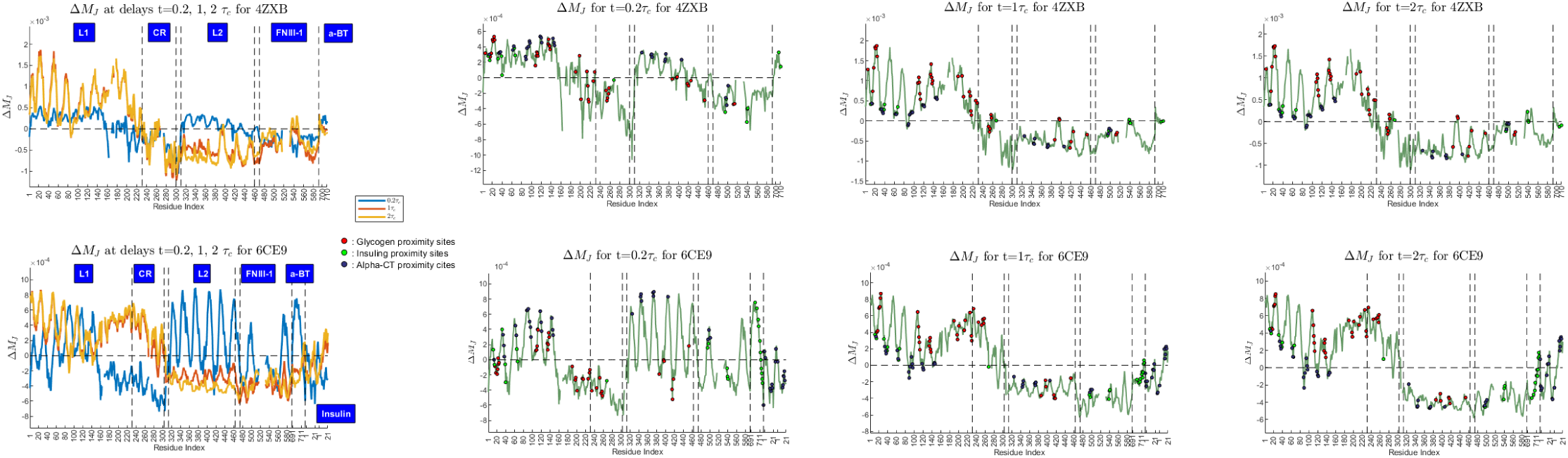
Δ*M*_*I*_ (*τ*) for the apo IR (4ZXB) at various time delays shown on only for the monomer. Filled spheres show functional residues (atomistic distance of which to the given ligand is less than or equal to 5*A*° see the legend box). The blue boxes and dashed vertical lines are used to depict different regions/domains. The narrow regions that are not labelled correspond to the linker 1 and linker 2 regions.

**Figure 7:**
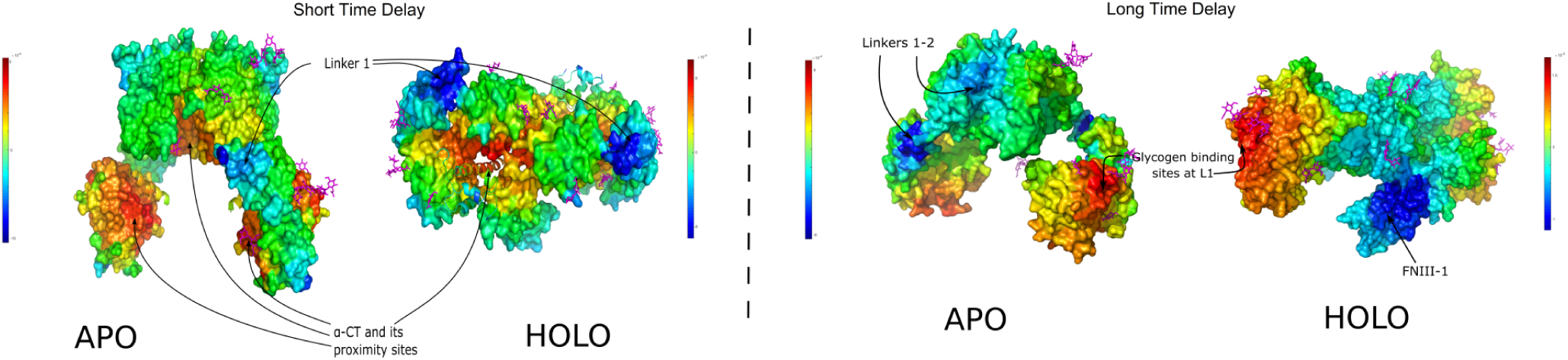
The apo (4ZXB) and holo (6CE9) states color-coded with Δ*M*_*I*_ (*τ*) values for *τ* =0.2 *τ*_*c*_ and 2 *τ*_*c*_ (negative values linearly to blue-cyan-green and positive values linearly to green-yellow-red color scales).

When the IR assumes the holo state, the positive peaks are at the *α*-CT itself (with the insulin binding sites) and the *α*-CT binding sites on the L1 and L2 domains. These peaks are much more emphasized compared to the positive peaks in the L1 and L2 domains of the apo state. We also see a few positive peaks at the FNIII-1 domain close to the *α*-CT proximity sites, which are not visible in the apo state. The positive peaks at the glycogen proximity sites do not persist. The negative peaks appear in the similar positions as in the apo state with an additional negative peak at the insulin molecule (which also includes some *α*-CT proximity sites).

Combining these observations with the 2D plots in Figure S3, the *α*-CT helix itself and the *α*-CT binding residues in the holo state are preceding and the linker 1 with the C-terminal of the L1 domain (the part that gets closer to FNIII domain in the holo state), the CR domain and the insulin molecule are lagging in their dynamics. This suggests a control of the *α*-CT helix and its binding sites on the IR mainly over the regions near the linker 1, the CR domain and the insulin molecule. This pattern is much more evident in the holo state, however a weaker form exists intrinsically in the apo state (see also Fig S4).

#### 3.2.2 Long time delay scale

In the apo state, all the highest positive peaks occur at the glycogen proximity sites in the L1 domain or their vicinity. The tallest negative peaks occur at the CR and L2 domains with the linkers 1 and 2 and some of the *α*-CT proximity sites, followed by some shorter negative peaks at the FnIII-1 domain. In the FNIII-1 domain, the *α*-CT proximity sites form some of the tallest negative peaks (Fig 6).

In the holo state, the tallest positive peaks still occur at the glycogen proximity sites of the L1 and CR domains. One of the most prominent changes is observed at the CR domain where the negative peaks turn into high positive peaks (some of which are at the glycogen proximity sites). On the other hand, some of the tallest negative peaks appear at the FnIII-1 domain, the linker 2 and the L2 domain followed by the *α*-CT. Except for the C-terminal end, the insulin molecule also displays negative peaks.

With 2D plots in Figure S3, the regions with high positive peaks preceding the dynamics of those with high negative peaks as mainly anticipated. There is also a pattern that is not evident in the 1D plots in Fig 6. His100-Glu180, which is a contact region between the L1 and FnIII-1 domains, while affected by the CR domain, has an influence on the dynamics of the FnIII-1 domain (particularly through Pro160-Glu180). An important distinction between the apo and holo states is the region corresponding to the L1 and FNIII-1 domains. We see that this region is red in the apo state and blue in the holo state which signifies that in the holo state the dynamics of the L1 domain precedes the dynamics of the FNIII-1 domain and in the apo state it is the converse.

Our findings overall suggest: In the holo state, the *α*-CT (and thus insulin proximity sites on it) and its proximity sites on the IR precede the behaviour of the rest of the IR, especially the linker 1 and the CR domain at the short time delay scale. In the apo state, even though less emphasized, this appears as an intrinsic remnant. This prompts us to suggest that the *α*-CT might be the key determinant in changing the conformation from the holo to other states. On the other hand, the *α*-CT also partially precedes the dynamics of the glycogen binding sites while affecting those on the CR domain but not those on the L1 domain. It is possible that anything that affects the stability of the interaction of *α*-CT with IR (such as dissociation of the insulin molecule from IR) induces a conformation change back to the apo state (see also Fig S4). It is likely that the facilitation of this pathway requires the *α*-CT to be in full contact with the IR to coordinate the dynamics of the rest of the protein.

We see that the glycogen proximity sites precede the dynamics of the rest of the protein for both apo and holo states in the long time delay scale. However, there are some marked changes between the two states. Only in the holo state, the glycogen proximity sites at the L1 and CR domain precede the dynamics of the FNIII-1 domain. For the apo state, the behavior is actually the opposite. Also, only in the holo state, the glycogen proximity sites precede the dynamics of the *α*-CT. These observations in the short and long time delays suggest the followings:

1. The conformational transition from the apo to holo states facilitated by the *α*-CT and insulin molecule at the short time scale brings the IR to a conformation where the glycogen proximity sites and CR can precede the dynamics of the FNIII-1 region, and thus possibly send signals downstream to the TK domain via this interaction.
2. Since the *α*-CT has a role in the conformational transition at short time delay scale, it is possible that glycolysation might also indirectly affect the conformational transition from the holo to the apo states by its preceding dynamics over the *α*-CT helix.

In fact, for some proteins, including the IR, glycosylation has been shown to affect the stability of a protein in a given state and thus possibly have a role in the conformation changes and function (see [18, 17, 16]). The glycogen proximity sites might thus act like a regulator on the function through the conformational change as we observe for the R domain of ATCase. To this end, our findings suggest that 1-glycosylation might affect the IR function through its interaction with the FNIII domain and the IR conformational stability via its interaction with the alpha-CT helix 2-glycosylation might affect the interaction with the TK domain via the FNIII domain once IR assumes the holo conformation.

### 3.3 DNA-dependent Protein Kinase (DNA-pk)

DNA-dependent Protein Kinase (DNA-pk) is a complex system formed by DNA-dependent protein kinase catalytic subunit (DNA-pkcs, 4128 residues) and Lupus KU Autoantigen proteins (KU70/80, 1341 residues) heterodimer, which together forms a tunnel to which the DNA binds. Major parts of the DNA-pkcs are the N-terminal domain, the circular cradle domain, the FAT domain and the kinase domain (important regions: See Fig 8 and Table S4). KU70/80+DNA binds to the DNA-pkc at the N-terminal and CC domains. The DNA-pkcs and KU70/80 has a scaffolding role in repairing DNA double stranded breaks through non-homologous end joining (NHEJ) or Variable Diversity Joining (VDJ) recombination or homologous recombination (HR). NHEJ and HR are competitive pathways and the realization of one is considered to block the other pathway. NHEJ starts when KU70/80 recognizes a broken DNA. Then, DNA-KU70/80 binds to the DNA-pkcs and acts as an allosteric regulator. This turns on the DNA-pkcs’s kinase activity. Even though the exact role of the kinase activity in NHEJ is unknown, it is known to be necessary for the phosphorylation of other NHEJ factors [27]. HR pathway is less understood, however for its activation, BRCA1 was proposed to bind to a site nearby the KU70/80 binding sites on the DNA-pkcs.

**Figure 8:**
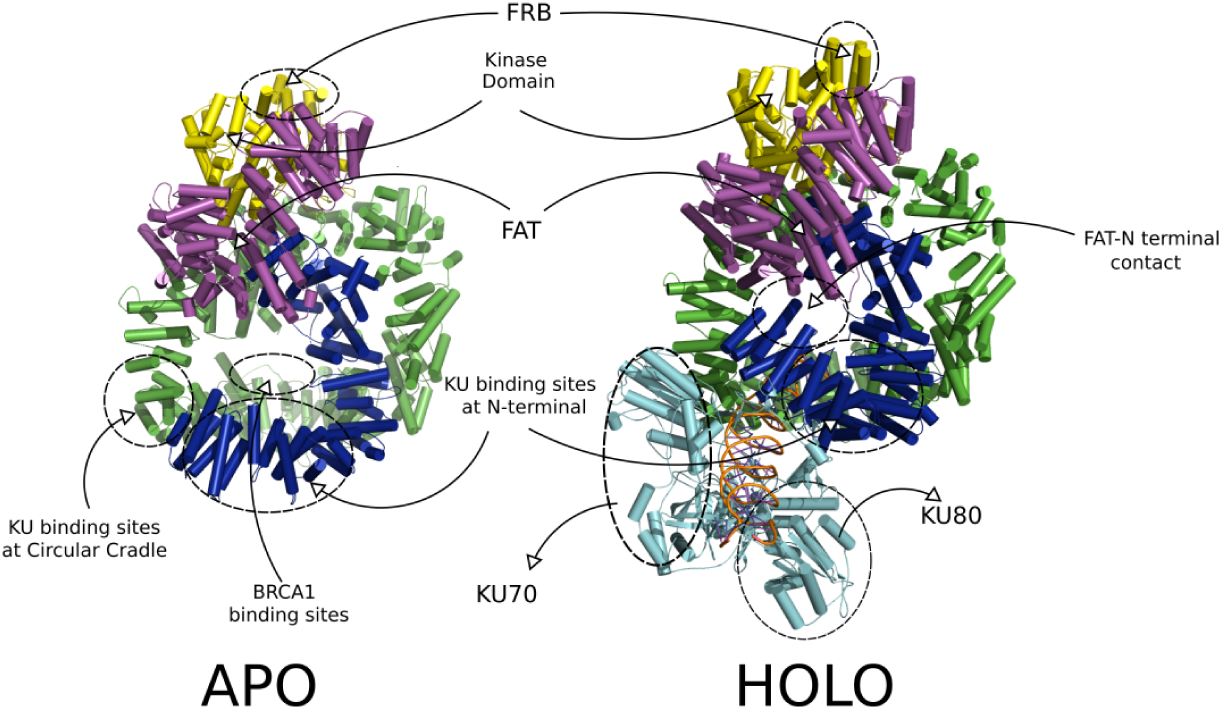
The apo (5LUQ) and holo (5Y3R) states of the DNA-pkcs. The holo state is in complex with the KU70/80 dimer (included in calculations) and DNA (DNA not included in both calculations). The major domains are colored as Dark blue: the N-terminal domain, Green: the CC domain, Purple: the FAT domain, Yellow: the KD, Sky blue: KU 70/80.

In the transition from the apo to holo states, major structural changes in the N terminal domain and minor changes in the FAT domain and the KD are realized. In particular, the first half of the N-terminal domain (up to residue 382), which is approximately 50 *A*° away from the FAT domain, rotates upwards and contacts the first half of the FAT domain (approximately up to residue 3314). This causes some rearrangements in the last half of the FAT domain and at the KD (especially at the FRB region). These rearrangements, by opening the catalytic gate, expose the catalytic groove leading to a wider entrance for substrates ([35, 27]). Residue 382 is called the switch point [35, 27], since the rest of the N-terminal is left essentially unchanged compared to the first half with global movements. The conformational differences of the apo and holo states suggest the movement mediated by a hinge region that connects the CC domain to the FAT domain and the KD [8]. Most of the computationally calculated hinge residues (global hinges for the apo and holo states were calculated using [9], listed in Table S4) belong to this region, backing this proposition and suggesting their importance in function.

There is yet no high-resolution structure for the apo/holo states of DNA-pkc. The available ones being are around 6 Å ([35, 27]) and the allosteric mechanism is still being debated. However, one proposed mechanism ([35, 27]) is that the DNA bound KU70/80 protein is required for the activation of the NHEJ pathway. There is no resolved structure of DNA-PKcs with BRCA1, however the underlying mechanism for the competition between NHEJ and HR pathway is believed to be allosteric and steric interactions between BRCA1 and the KU70/80 protein. There is also experimental evidence that the DNA-pkc may be activated in a DNA independent manner through C1D that binds at the Leucine zipper motif (1503-1538) in the CC domain [36].

In this case, we carried out DORIA on the apo (5LUQ) and holo (5Y3R) states. The apo state doesn’t contain the KU70/80 heterodimer, but the holo state does.

#### 3.3.1 Short to Normal Time Delay Scales

##### The apo state

The time delay scales *τ* =0.4 *τ*_*c*_ to 2 *τ*_*c*_ show qualitatively similar behavior, however the relative height of the peaks change from one to the other (Fig 9, see also Figure S5 for 2D). At *τ* = 2*τ*_*c*_ time delay scale, the highest positive peaks occur at residues 506-1120 that contains the phosphorylation cluster JK and HCR I followed by other peaks at HCRIII and HCRIV, V (respectively in the FAT domain and the KD). We approximately call the union of these regions as the Central Regulatory Region (CRR) for the apo state. It is interesting to note that this region also includes the global hinges that appear both in the apo and holo states. Note that even though the places where the positive peaks occur are far in sequence number, spatially they are closely located towards the center of the DNA-pkcs, hence also justifying the name CRR (see also Fig 11).

**Figure 9:**
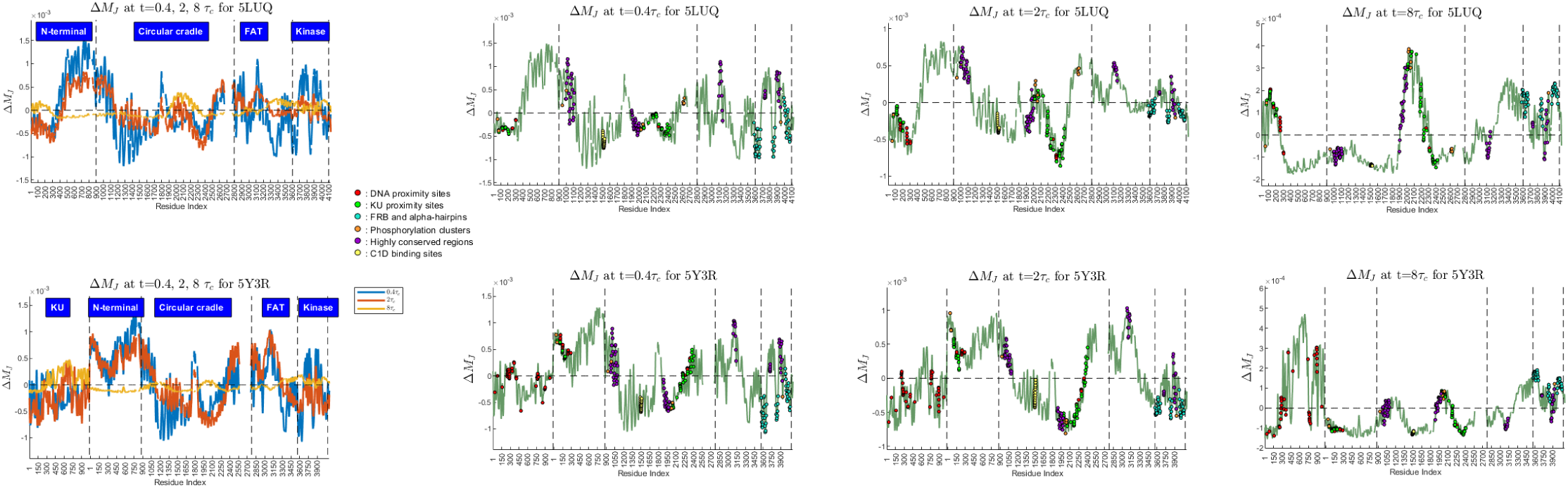
Δ*M*_*J*_ for 5LUQ and 5Y3R at the short, normal and long delay scales. The disks for different phosphorylation clusters are all colored with the same orange color. BRCA1 is assumed to be around PQR phosphorylation cluster (which is near residues 2020-2050) and not shown separately.

**Figure 10:**
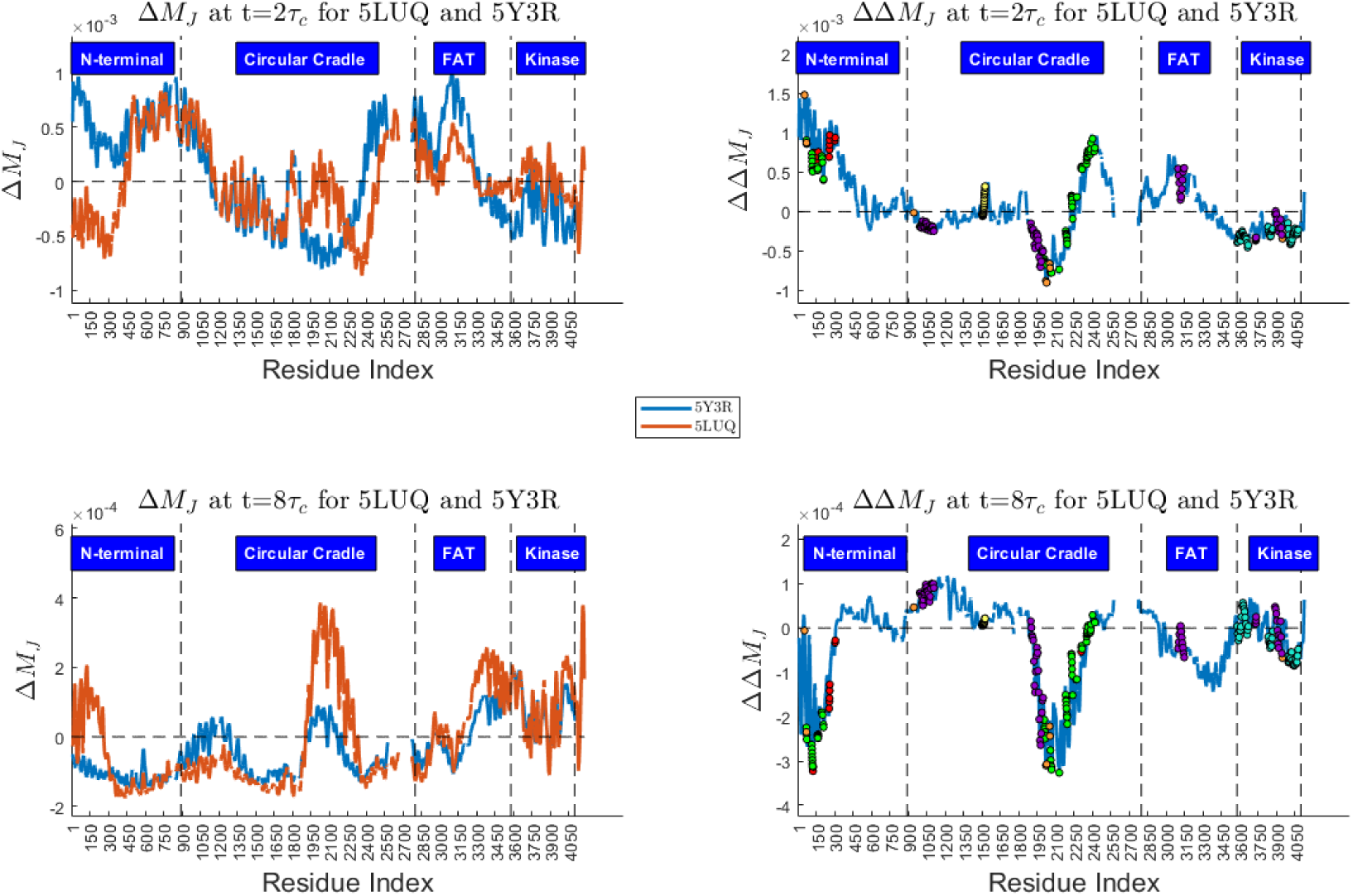
Comparison of Δ*M*_*J*_ (*τ*) for the apo and holo states

**Figure 11:**
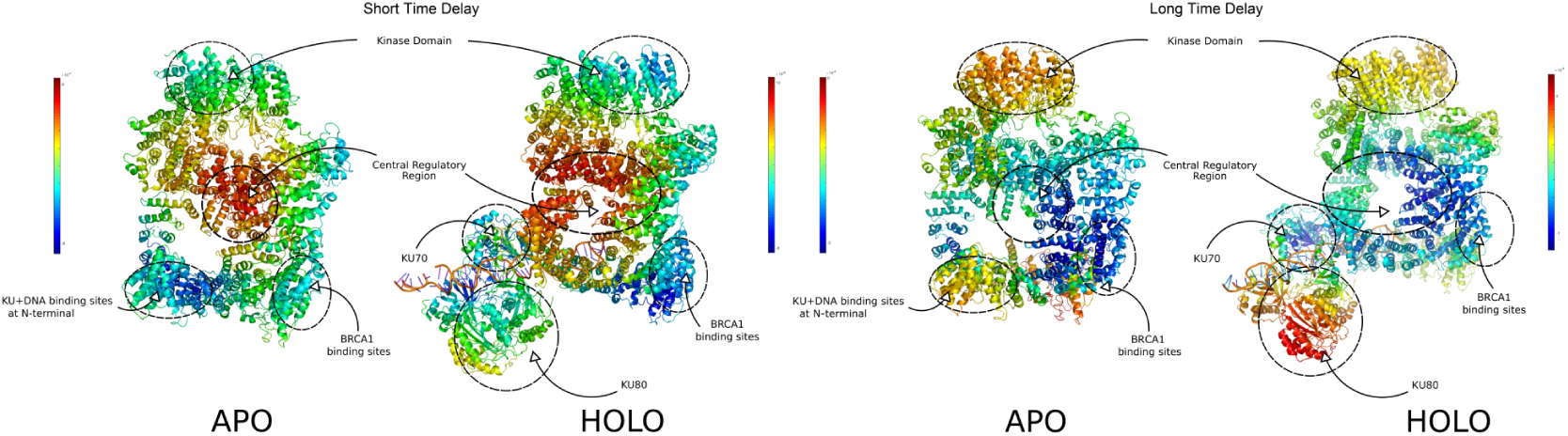
The Apo (5LUQ) and holo (5Y3R) states colored according to values of Δ*M*_*I*_ (*τ*) t=0.4 and 8*τ*_*c*_.

The highest negative peaks occur at the KU binding sites in the CC and N-terminal domains, some regions of the CC domain (roughly residues 1200-1800) that contains HCRII and a global hinge (residue 1421), the C1D binding sites, the PQR phosphorylation cluster, the BRCA1 binding sites, the catalytic gate in the KD and a region of the FAT domain (around residue 3376). We also see that the main differences between 0.4*τ*_*c*_ and 2*τ*_*c*_ is that the negative peaks at the first half of the CC domain and the catalytic gate become shorter whereas others become taller for 2*τ*_*c*_ with similar patterns.

2D plot reveals (Fig S5) that the CRR region is preceding and the regions with the negative peaks lagging in their correlated dynamics, suggesting the controlling role of the CRR in this delay window. The residues 1200-1800 of the CC domain even though has an overall negative peak in 1D plots (Fig 9), 2D plots reveal that it is not always lagging but also has preceding fluctuations at some regions. In fact, preceding and lagging regions respectively correspond to the KU binding sites and the CCR domain.

##### The holo state

Fig 9 ad Fig 10 reveal the changes with the conformational change from the apo to holo states at *τ* =2 *τ*_*c*_. The most prominent change is that the KU binding sites in the N-terminal domain switches from negative to positive values. The double negative peaks at the KU binding sites and HCRII in the CC domain merge to a single positive peak at the BRCA1 binding sites. The height and width of the positive peak at the FAT domain increases (especially at HCRIII). Like the apo state, there is a central regulatory region (CRR) -larger in size compared to the apo state- including the KU binding sites in the N-terminal domain as well as the region of the FAT domain around HCRIII. There is a general increase of the negative peaks at the KD (concentrated more on the catalytic gate and partially on HCRIV).

2D plots (Fig S5) display the preceding behavior of the CRR and the lagging behavior of the KD as in the apo state, suggesting its control role on the catalytic gate. However, unlike the apo state, this control persists up to longer delay scales of 2*τ*_*c*_. One can look at these 2D plots to see which interactions contribute to the increased control on the catalytic gates at the short time scale when the protein switches to the holo state. Looking at the column corresponding to the N-terminal of the KD; the most blue regions (i.e regions that control this gate) correspond to the second half of the N-terminal unit in the apo state, whereas the intensity of the blue color in the first half of the N-terminal unit and the FAT domain increases to the same level in the holo state. This confirms that the FAT domain and the region of the N-terminal unit become a part of the CRR in the holo state, which allows them to augment the control on the catalytic gates. This agrees with the predictions done in [35, 27].

#### 3.3.2 Long Time Delay Scale

##### The apo state

The highest positive peaks occur at the CC domain, concentrated in the region containing the BRCA1 binding sites, HCRII and the KU proximity sites. This is followed by smaller positive peaks at the KU binding sites in the N-terminal domain, the regions of the FAT domain close to the KD (residues 3350-3600) and some parts of the KD itself including the catalytic gate. Highest negative peaks appear at the CRR region (the hinge region in the N-terminal domain and the CD1 and the KU binding sites in the CC domain) and some of the KU binding sites in the CC domain.

2D plots (Figure S5) reveals that in terms of inferred causality the regions with high positive peaks listed above are preceding and the regions with the highest negative peaks lag in their dynamics.

##### The holo state

With the conformational changes from the apo to holo states, the highest positive peaks occur at KU80 (especially at residues His152-Met210). The rest of the positive peaks at the CC domain, the FAT domain and the KD appear as a suppressed version of the positive peaks of the apo case. The tallest negative peaks occur at the CRR, the C1D binding sites, the KU binding sites (both in the CC and N-terminal domains). Finally first 150 residues of KU70 that includes some of the DNA proximity sites and the sites at which KU70 interacts with the DNA-pkc and the DNA proximity sites on KU80. 2D plot (Figure S5) suggests that the KU strongly precedes the dynamics of the most of the protein, i.e. suggesting a controlling role for the KU proteins. The other regions including also the KD, which are positive but shorter than the KU, are controlled by the KU but precede the dynamics of the rest of the protein. The effects of these regions are more strongly present in the apo state.

An interesting change from the apo to the holo state in the long time delay scale is that the BRCA1 binding sites’ effect on the dynamics of the rest of the protein is almost completely suppressed in the holo state and the positive peaks at the KU binding sites in the N-terminal domain of the apo state become negative in the holo state.

It is worth to note that when the protein changes its conformation from the apo to holo state, one also observes a change in some of the global hinges [9]. In particular, a new hinge at residue 96 of the N-terminal domain appears and the hinge at residue 855 of the FAT domain shifts towards residue 848, which is closer to the N-terminal domain. This is inline with the observation that in the holo state the point of contact between the N-terminal unit and FAT domain starts to have influence on the dynamics of the protein. This likely creates a cluster of residues that has enhanced effect on the dynamics of the KD, stronger than the isolated effect of the FAT domain on the KD of the apo state.

##### Overall, the findings imply

1. We propose the existence of a central regulatory region CRR acting via global-dynamic interactions or in other words by the cooperative behavior of large rigid dynamic domains at the shorter time delay scales. Both in the apo and holo states, the CRR roughly corresponds to a region between the lower half (the N-terminal and CC domains) and the upper part (the FAT domain + the KD) of DNA-pkc. More precisely, in the apo state, the CRR is clustered around global hinges and HCRs I, III, IV, V. The CRR precedes the dynamics of the KU/DNA proximity sites in the N-terminal, the KU proximity sites in the CC domain, some regions around the C1D binding sites and to some extent the KD. In the holo state, however, when the N-terminal contacts the FAT domain, not only the N-terminal (including the KU/DNA proximity sites there) domain starts to regulate other regions but also the influence of HCRIII (inside the FAT domain) on the rest of the protein is enhanced. In particular, the effect of HCRIII on the catalytic gate region of the KD (FRB) is enhanced up to the time delay scales 2*τ*_*c*_. This appears to be due to the change in the contact numbers between the N terminal and FAT domains. This supports the suggestions put forward that when the KU/DNA complex binds to the DNA-pkc, it allosterically regulates the catalytic activity by perturbing the residues forming the gate near the catalytic cavity and the allosteric signal is transmitted via the FAT domain [35, 27]. It is also worth noting that the CRR precedes the dynamics of the KU and it is thus also possible that it regulates the disassociation of the KU from the DNA-pkc via global-dynamic interactions.
2. It has already been suggested that KU80 mediates an allosteric activation of the DNA-pkc for the NHEJ pathway [28]. One of the proposed mechanisms was that KU80 partially blocks the BRCA1 binding site thus favoring the NHEJ pathway over HR. We extend this proposition by the following observations: The region His152-Met210 in KU80 might have several regulatory roles via sub-domain dynamic interactions. At longer time delay scale, this region precedes the dynamics of the binding sites between KU70 and the DNA-pkc, the CRR and the BRCA1 binding sites. KU80 might thus have an allosteric regulatory role (apart from the possible steric clash) over the BRCA1 and the KU binding. It might be worth studying whether if KU80 has some potential ligand binding sites that might act as regulators. Since this region also precedes the CRR (which contains the hinges), it is also possible that regulatory molecules attaching to this region might allosterically affect the function and conformational preference of the DNA-pkc as we have seen in the regulatory region of ATCase.
3. At long time delay scale, in the apo state, the KU and BRCA1 proximity sites and the KD (and the last half of FAT domain close to it) precede the dynamics of the rest of the protein including the CRR region suggesting their possible allosteric control over the protein’s dynamics. Upon the KU binding, the control almost entirely passes over to the KU80 protein (see item 2 above). This suggests two things; a- Regulatory molecules/proteins (such as KU) binding to these regions have allosteric effects over the rest of the protein (which is also inherent but weaker in the apo form), b-The binding of regulatory molecules to one of these regions may have a negative allosteric effect on other remaining regions preventing regulatory molecules from binding there.

## 4 Conclusion

In this paper we put a dynamic order on residue fluctuations by using the asymmetry in mutual information *M*_*IJ*_ (*τ*) (which is completely determined by *C*_*IJ*_ (*τ*) in a straight forward manner). By dropping the high friction assumption we are able to do this in the regime of ENMs and apply it to very large proteins easily. By studying a text book example, ATCase, we demonstrate the relations between this dynamical order, function and allosteric behaviour. Dynamical order for short delay scales is related to global dynamics of the protein with large rigid body movements where as for long delay scales more intricate allosteric behaviours probably involving more subdynamic domains are observed. In the light of this we apply this to IR and DNA-pkc to both test known hyphotheses about these proteins as well as pose new ones. The formulation of this model using stochastic differential equations makes it very transperant to modifications such as studying triple interactions rather than pairwise or looking at perturbative effects of changing bond structure to dynamical order. We will explore these directions further in upcoming works.

## Acknowledgements

All figures were created using PyMOL and Inkscape. The initial parts of the paper was completed when the author was at Imperial College London where he acknowledges partial support by ERC AdG grant no: 339523 RGDD. The second author acknowledges support by TUBITAK (The Scientific and Technological Research Council of Turkey) grant no: 115M418.

## Code Availability

The custom code written in MATLAB which is used to analyze the proteins in the manuscript is available upon request.

## Data Availability

The data representing the delayed communication profile at chosen delay times for each protein is available upon request.

## Contributions

S.T formulated the mathematical method and programmed the code for application of the model. Both authors contributed to conceptualization of the project, study of examples in the article, interpretation of results and preparation of the manuscript. T.H initiated and supervised the project.

